# Habitat suitability modeling of Himalayan Monal and Koklass Pheasant in Western Himalayas and Hindukush, Pakistan

**DOI:** 10.1101/2022.08.17.504340

**Authors:** Muhammad AzharJameel, Muhammad Sajid Nadeem, Muhammad Kabir, Tariq Mahmood, Faraz Akrim, Muazzam Ali Khan, Muhammad Naeem Awan, Muhammad Fiaz Khan, Muhammad Zubair Anjum, Shahzad Aslam

## Abstract

The Himalayan pheasants are under the greatest threat due to habitat degradation, and loss. Quantifying geographical range and suitable habitat of a species can help in better management and conservation decisions. Himalayan Monal *(Lophophorus impejanus)* and Koklass *(Pucrasia macrolopha)* are endemic to the Himalayas and Hindukush mountains. This study aims to investigate habitat suitability of these pheasants in the western Himalayas and Hindukush. MaxEnt and Cringing models were used to document habitat suitability and to identify valleys with most suitable habitat. MaxEnt model displayed excellent predictive performance showing a strong prediction of the probability distribution and habitat. The area under cover (AUC) values quantified for the replicate runs were 0.994 (±0.001) and 0.991 (±0.005) for Himalayan Monal and Koklass pheasant respectively. The climatic parameters including temperature, precipitation of the warmest quarter (bio_18) contributed the maximum 21.3% and 23.5%, followed by annual precipitation (bio_12) 12.3% and 8.9% for habitat prediction of Monal and Koklass. The topographical variables, altitude, slope, and distance to settlements contributed 15.2%, 2.6%, and 16% in the Monal habitat prediction model while 8.4%, 10.5%, and 15.8% for the Koklass habitat prediction model respectively. We quantified highly suitable (844.4 sq. km), moderately suitable (2819.42 sq. km), and less suitable (3933.09 sq. km) habitat for Monal pheasant. Whereas, highly suitable habitat for Koklass pheasant was (611.5 sq. km), followed by moderately suitable (2551.3 sq. km), and less suitable (4494.11 sq. km). Bar Palas region of Koli Palas district, Jalkot and Kandia valley of district upper Kohistan and Kayal valley of district lower Kohistan were identified as core zones or hot spots for these pheasant species. Areas identified as core zone/hotspot and suitable habitat for the pheasant species should be legally protected for the conservation of pheasants.

## Introduction

The changes in environment affect both the ecology and habitat of many species around the world. The ecosystem of the Himalayas generally explicit such progressions, experiencing uncertainty on temperatures, precipitation, and water sources [1,2] Severe climatic and environmental fluctuations are evident in winter and summer [1] and the impact of these fluctuations vary from species to species. An understanding of the influence of these changes on the interaction of species is vital for their conservation and management [3]. The mountain ecosystems of the eastern Himalayas are poorly explored due to tough terrains and harsh climatic conditions. The climatic factor directly effecting the Himalayan endemic bird species influenced on dynamic, shifting and shrinkage of the habitats [4].

Species distribution models (SDMs) play an important role in the conservation and management of wildlife species and are used to gain ecological and evolutionary insights and help to predict current, and future distribution ranges of species of interest. SDMs have been helpful in restoring the threatened species that have been extinct in the regions [5,6]. The SDMs mostly used in hard terrains and the researchers generally used this method for the study of threatened species in hard environmental conditions and tough terrains [7]. The bioclimatic model is used to predict current and future suitable habitats for a species by using bioclimatic factors of the area of interest [8]. Maximum entropy modeling (MaxEnt) helps to develop a feasible geographic distribution model by using occurrence records and bioclimatic variables [1,9].

The Galliformes are a group of terrestrial birds, widely distributed in a variety of habitats including deserts, cultivated lands, forests, and alpine meadows [10]. The Himalayas have the highest diversity of Galliformes (n=34), out of which 29 species are range-restricted to the Himalayas and 25% of these species being threatened [11,12]. The populations of Himalayan pheasants are highly fragmented and declining [1]. Among the Himalayan range-restricted species, scientific data on 15 species is lacking due to shy nature and remote habitats. The pheasant species are threatened due to habitat loss, anthropogenic pressure, and global warming [13]. The Himalayan Monal *(Lophophorus impejanus)* and Koklass *(Pucrasia macrolopha)* are noteworthy species of pheasants distributed in the Himalayas and Hindukush ranges of northern Pakistan. These species are adversely effected by climate changes their geographical ranges shrinks and will shift toward higher altitudes in near future. These species inhabit the same macro habitat but utilize different microhabitat and altitudinal ranges. Their distribution has been reported in the northwest part of the western and eastern Himalayas, the lesser Himalayas and their foothills [10]. Himalayan Monal is known to occur in Pakistan, India, Bhutan, and Assam [14,15]. It inhabits the upper temperate forests of conifers, and broad-leaved forests with densely vegetated slopes at altitudes ranging from 2400 m to 4500m [16,17]. Koklass is confined to tree lines not extending beyond this range [18]. Koklass can reveal their presence by loud chorus calls during breeding seasons and autumn [19]. Considering the importance and range restrictions of these species, this study aims to, (1) investigate habitat suitability of Himalayan Monal and Kolass pheasant in the western Himalayas and Hindukush, with special reference to Indus Kohistan, (2) identify the core/hot spot zones for these species in the study area. Our study provides baseline information from the unexplored region of Pakistan vital for effective management and conservation of both pheasants in the study area.

## Methods

### Study area

The study was carried out in the western Himalayas and Hindukush ranges intensively focused on Indus Kohistan, Pakistan (N35º6’0’’-35º42’0’’, E72º36’0’’-73º48’0’’). Indus Kohistan is administratively divided into three districts upper Kohistan, lower Kohistan, and Koli Palas Kohistan [20]. Indus Kohistan is adjoining to Azad Kashmir from the east, district Swat in the west, Diamer (Gilgit Baltistan) at the north, Battagram, and Shangla from the south (Fig 1). The topographical condition of the area is a hard-rocky terrain with rugged undulating steep mountains. The climate of the area varies with monsoon-influenced moist temperate zones in the western Himalayas and semi-arid cold deserts in Hindukush, Pakistan [21]. The Indus Kohistan region of the Himalayas consists of moist temperate evergreen oak and conifer forest [22]. This region is enriched in resources including dense forests, extensive grasses, and arable lands for mineral deposits [22]. The local people experience seasonal migration (summer and winter migration) with their livestock. Pheasants species recorded from the study area include Chukar *(Alectoris chukar)*, Himalayan Monal *(Lophophorus impejanus)*, Koklass pheasant *(Pucrasia macrolopha)*, Western Horned Tragopan *(Tragopan melanocephalus)* Himalayan snowcock *(Tetraogallus himalayensis)* [23]. The study area has great potential for pheasants and is one of the significant habitat for pheasant breeding and inhabits the world’s largest population of western horned Tragopan [24].

**Fig 1.**
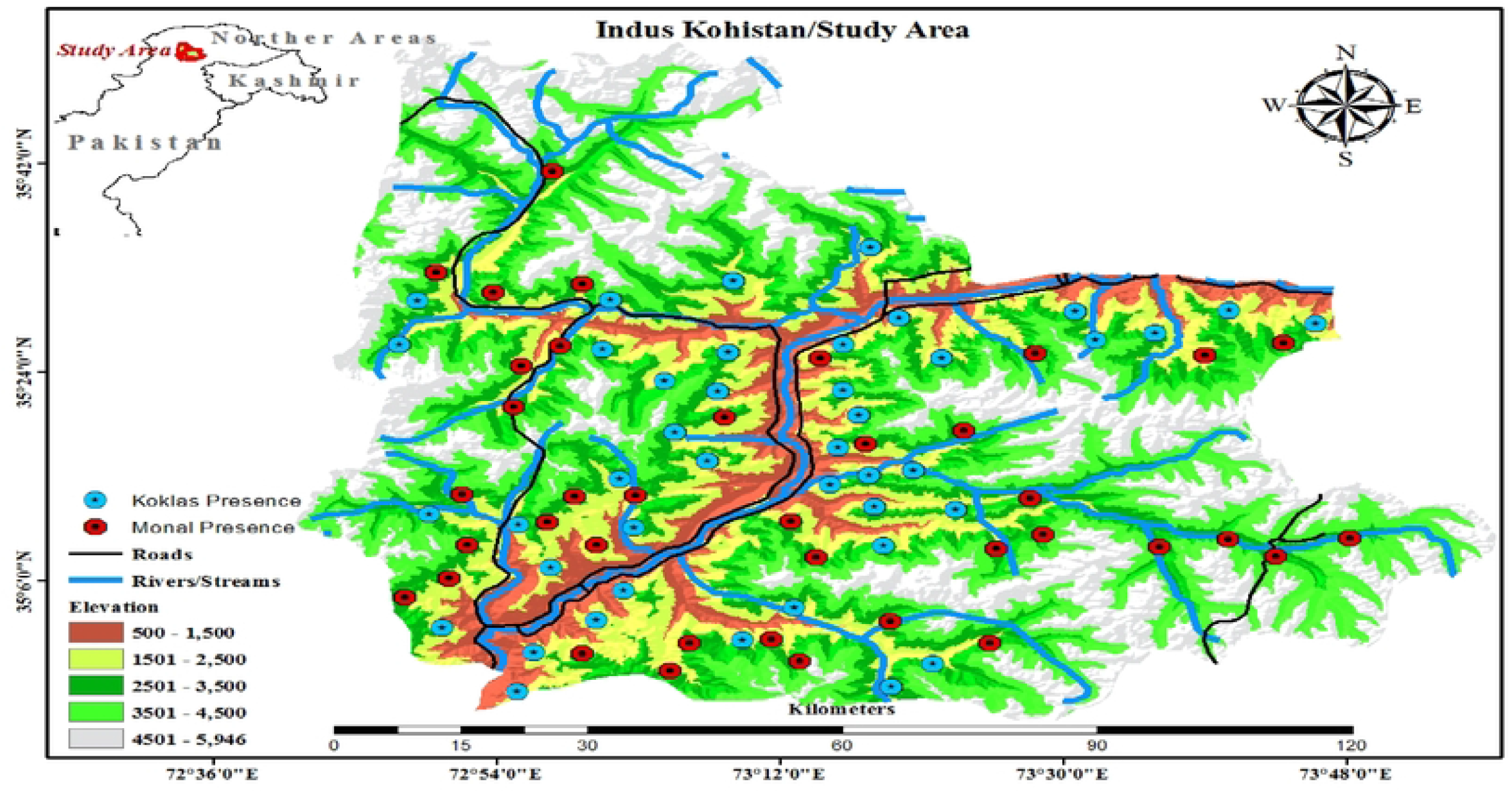
Digital Elevation Model of the Study area with Presence Points of Pheasants

### Study design

The duration of the study was from 2019 to 2021 and data were collected from six valleys of the whole study area. These valleys covers most of the region of the study area, selected on the basis of previous literature, visual assessment in pilot project, and less human interference. Three valleys Palas, Jalkot, and Harban were selected from the western Himalayan range, and three valleys; Dubair, Kayal, and Kandia were selected from the Hindukush range respectively. The total of 36 surveys covered 110 vantage points preferably in summer (March-June) in the breeding season of pheasants. We have collected a few occurrence data in winter from lower altitudinal vantage points specifically lower altitudinal species. The pheasants generally call in early dawn and dusk (Just before sun rise and after sunset) and we reached the vantage point accordingly. Each vantage point was surveyed twice to confirm the occurrence and effort lasted for 60 minutes in each specified point.

### Presence record

The occurrence data of Himalayan Monal and Koklass pheasant were collected by using the primary data collection method (feather collection, call records, flush method/direct observations), these methods are widely used in hard terrains and remote areas for population and occurrence recording of the pheasants [1] and also confirmed the presence by secondary data from the local community of potential pheasant habitats [24]. A total of 68 and 51 occurrence points of Himalayan Monal and Koklass pheasant were recorded respectively from the study area. We used 2Km spatial autocorrelation to remove spatially correlated data points and generated independent locations. After the selections from an initial dataset of the occurrence records, only 55 and 42 unrelated deviated locations of Himalayan Monal and Koklass pheasant were used respectively to generate the current models (Table 1) [25].

**Table 1.** Himalayan Monal and Koklass Pheasant Presence Points in all the valleys of the study area.

| S. No | Valleys | Himalayan Monal |  |  | Koklass Pheasant |  |  |
| --- | --- | --- | --- | --- | --- | --- | --- |
|  |  | Direct Observation | Call Records | Feather collection | Direct Observation | Call Records | Feather collection |
| 1 | Palas Valley | 3 | 3 | 4 | 2 | 3 | 4 |
| 2 | Jalkot Valley | 7 | 5 | 14 | 5 | 6 | 2 |
| 3 | Harban valley | 3 | 1 | 1 | 4 | 2 | 0 |
| 4 | Kandia Valley | 6 | 4 | 7 | 6 | 8 | 2 |
| 5 | Kayal Valley | 3 | 2 | 1 | 3 | 2 | 0 |
| 6 | Dubair valley | 2 | 2 | 0 | 1 | 3 | 0 |
| Total presence points |  | **68** |  |  | **54** |  |  |

### Variables selections

To generate the model, we consider initially a set of 28 environmental variables including, altitudes, 19 bioclimatic variables, land cover, slope, soil type, distance to road, distance from rivers and streams, and distance to human settlements, vector ruggedness measure, and normalized difference vegetation index. Bioclimatic variables and altitudes were obtained from the worldClim database (www.worldclim.org/current) [26]. Land cover was obtained from Global land cover available from (https://lta.cr.usgs.gov/glcc/globdoc2_0) [21]. Distance to roads, rivers, human settlements was analyzed by using Euclidean distance tool in Arc GIS 10.5. Soil (FAO, 2003, digital soil map of the world), vector ruggedness measure (SRTM 90m DEM by Center for Nature and Society, Peking University) vegetation index, and its normalization were obtained from the NASA website (http://modis-land.gsfc.nasa.gov/vi.html). The finalized eight retained variables used for the model were; soil, altitudes (m), slopes, global land cover 2000, settlement distance (km), mean NDVI, temperature (ºC), and precipitation (mm).

### Model execution

To remove the variables which are highly correlated before generating the models, we defined the correlation matrix by using Pearson’s techniques and selected the only variables with a correlation of less than 70% (r < 0.70) [27]. We selected the set of variables that were prime representative of the pheasant’s ecological requirements in the specific area [1]. After the analysis eighteen variables were finalized for their applicability to the scale of this study [28]. All the variables were prepared conforming cell size [30-arc second resolution (0.93 × 0.93 km = 0.86 km^2^ at the equator)], geographic extent, projection, and ASCII format using the ‘resample’, ‘clip’, ‘mask’, and ‘conversion’ tools in Arc GIS 10.5.

### Maxent model and its validation

MaxEnt (version 3.2.4) was used for habitat suitability modeling of two pheasant species [9]. When absence data has high uncertainty, then the recommendation is to use presence which is especially true when detection rates are poor [21]. We used the Maxent model for the distribution of both pheasant species and recognized it as better performing with presence-only data, especially with the minimum number of occurrence records. For model preparation, we used the presence record defined as “Monal/Koklass occurrence point” respectively, and the environmental variables defined as “environmental layers” in Maxent. The setting panel of the maxent we selected auto features; random seed; write plot data; remove duplicates presence records; give visual warming; show tooltips; regularization multiplier; maximum 10,000 background points; iteration maximum 1,000. Finally, we archived a five replicates effects with cross-validation of Pearson’s techniques. These settings are conservative enough to optimize the performance of the model and get close to convergence [9]. The percentile training presence like 90% of the training locations classified correctly was selected for the threshold value to show the presence of the species, and also reclassify our models into binary presence/ absence map. This is a conservative method commonly adopted for species distribution modeling, particularly the studies which rely on datasets and data collected by using different observers and methods [29]. Furthermore, a Jackknife test was performed to select the requisite bioclimatic variables, which may differ from the precedent variables. In general, three plots are accustomed to showing variable gravity, named as regularized terrain gain, test gain, and AUC of the test data. The Jackknife-cross-evaluation test revealed the relative contribution and permuted by each effective environmental variable through Maxent. The whole maxent process generated three models; first, the model was created by using the variables to check which variable is most informative. Second, a model is created individually by using each variable to find which had the most information not featured in the other environmental variables. The final model was generated based on all variables and response curves derived from the univariate model were plotted to predict the single environmental variable influence presence probability.

We tested the model by using different validation methods; the receiver operated characteristics, analyzing the area under the curve (AUC), and true skill statistics (TSS). AUC accesses the discrimination ability of the models and values ranges from 0 (equaling random distribution) to 1 (perfect prediction). The value of AUC >0.75 corresponds to high discrimination performance. TSS compares the data as correct forecast number minus random guessing, to that of a hypothetical set of perfect forecasts. That considers both omission and commission errors and success as the result of random guessing; the values ranging from -1 to +1 correspond to perfect agreements and 0 or less to a performance no better than random [30].

## Results

### Habitat suitability

The Maxent model perfectly predicted that there is a suitable Himalayan Monal and Koklass pheasant habitat in the study area and their adjacent valleys in other districts of Khyber Pakhtunkhwa. The suitable fragmented habitats were predicted in Jalkot, Kandia, and Palas valley by the model, and some suitable habitat patches were also predicted from Dubair, Kayal, and Harban valley. These areas are mostly less altered by human interference, significant forest cover, and soft terrains with gentle slopes. The model also predicted that the valleys which have more human settlements occupied the microhabitat of pheasants, and open land covers were predicted as less suitable for the pheasants.

The MaxEnt displayed overall “excellent” predictive performance showing a strong prediction of the probability distribution. The regularization multiplier chose at 0.1 because of its ideal for the model sensitivity and area under the curve (AUC) against other multipliers. The AUC values quantified for the average replicate runs were 0.994 (±0.001) and 0.991 (±0.005) for Himalayan Monal and Koklass pheasant respectively. Among the environmental parameters, temperature like precipitation of the warmest quarter (bio_18) contributed the maximum of 21.3% and 23.5% for habitat prediction of Monal and Koklass, followed by annual precipitation (bio_12) 12.3% and 8.9% respectively. The topographical variables, altitude, slope, and distance to settlements contributed 15.2%, 2.6%, and 16% in the Monal habitat prediction model while 8.4%, 10.5%, and 15.8% for the Koklass habitat prediction model respectively. The biophysical (NDVI) have no such contribution for these species modeling in the study area while mosaic land cover (glc2000) contributed 11.1% and 12.9% for Monal and Koklass respectively (Table 2).

**Table 2.**
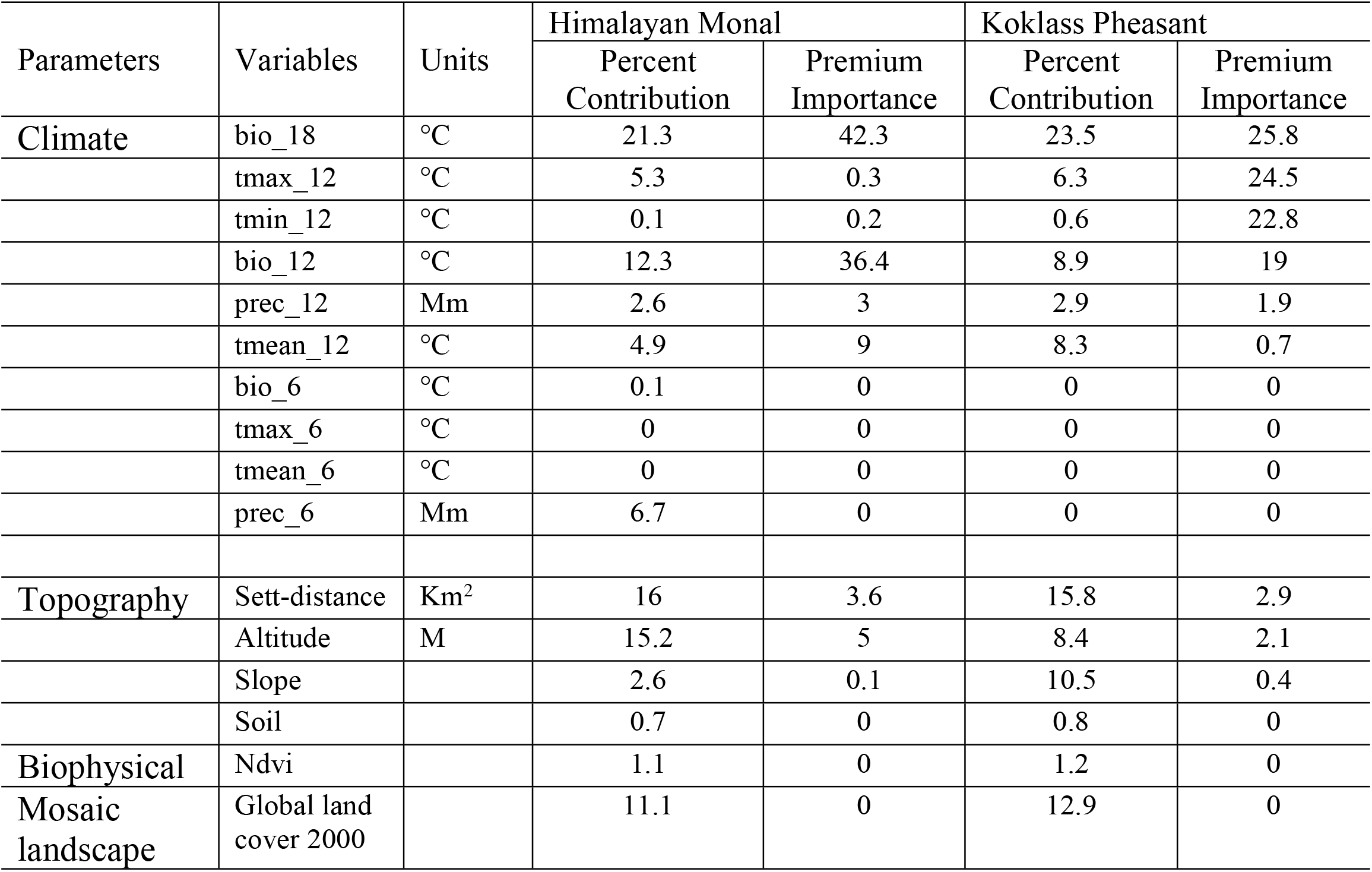
Environmental parameters and their contribution to habitat prediction of Monal and Koklass pheasant

| Parameters | Variables | Units | Himalayan Monal |  | Koklass Pheasant |  |
| --- | --- | --- | --- | --- | --- | --- |
|  |  |  | Percent Contribution | Premium Importance | Percent Contribution | Premium Importance |
| Climate | bio_18 | °C | 21.3 | 42.3 | 23.5 | 25.8 |
|  | tmax_12 | °C | 5.3 | 0.3 | 6.3 | 24.5 |
|  | tmin_12 | °C | 0.1 | 0.2 | 0.6 | 22.8 |
|  | bio_12 | °C | 12.3 | 36.4 | 8.9 | 19 |
|  | prec_12 | Mm | 2.6 | 3 | 2.9 | 1.9 |
|  | tmean_12 | °C | 4.9 | 9 | 8.3 | 0.7 |
|  | bio_6 | °C | 0.1 | 0 | 0 | 0 |
|  | tmax_6 | °C | 0 | 0 | 0 | 0 |
|  | tmean_6 | °C | 0 | 0 | 0 | 0 |
|  | prec_6 | Mm | 6.7 | 0 | 0 | 0 |
| Topography | Sett-distance | Km<sup>2</sup> | 16 | 3.6 | 15.8 | 2.9 |
|  | Altitude | M | 15.2 | 5 | 8.4 | 2.1 |
|  | Slope |  | 2.6 | 0.1 | 10.5 | 0.4 |
|  | Soil |  | 0.7 | 0 | 0.8 | 0 |
| Biophysical | Ndvi |  | 1.1 | 0 | 1.2 | 0 |
| Mosaic landscape | Global land cover 2000 |  | 11.1 | 0 | 12.9 | 0 |

The Jackknife regularized training gain variable importance for Monal and Koklass highest gain when used in isolation, average annual mean temperature (tmean_12) and minimum temperature of the coldest month (bio_6) differed the contribution for habitat suitability prediction (Fig 2).

**Fig 2.**
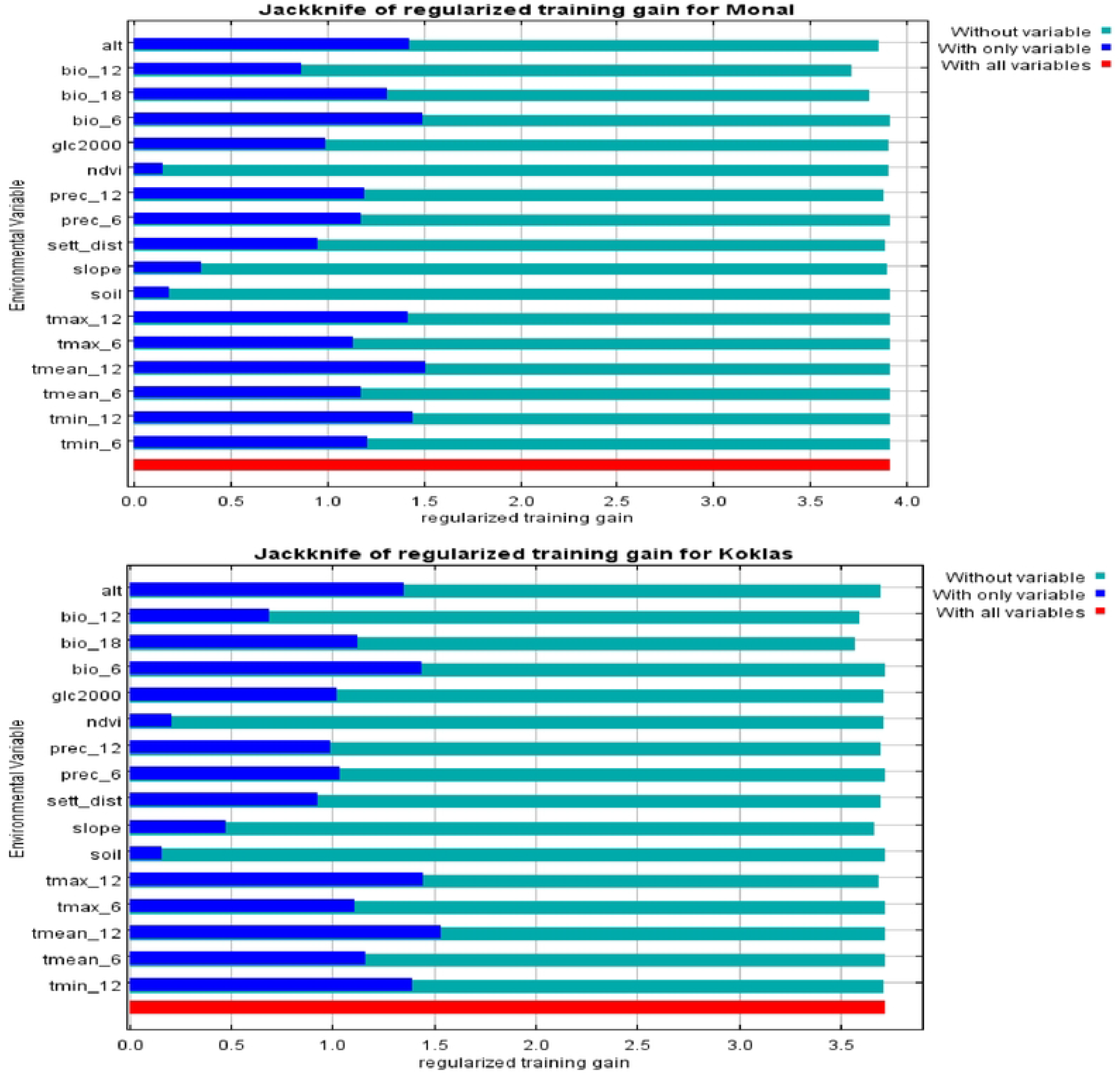
Jackknife analysis of environmental variables for Himalayan Monal and Koklass pheasant: It shows the importance of each variable in explaining the Pheasant species’ presence when used separately (cobalt blue), and how the model is affected when each variable is left out (aqua). Dark blue bars = Importance of single variable, light blue bars = loss in model gain, when the variable is omitted. Red bar = total model gain. Alt = Altitude, ndvi=normalized division of vegetation index, minimum temperature of coldest month (bio_6), annual precipitation (bio_l2), average annual mean temperature (tmean_12), precipitation of warmest quarter (bio_18), precipitation (1-12), maximum temperature (max6-12), minimum temperature (min6-12), average temperature biannually (mean-6), global land cover (glc2000) distance to settlement (sett_distancc), slope (slope), and soil (soil).

### Suitable area and hot spots

The current study quantifies the area of the suitable habitat and core zones of the pheasant distribution. Geographical distribution of pheasants predicted by the MaxEnt model occupying the ranges from the lower regions of a sub-tropical thick forest of oaks, tropical conifer forest including deodar, fir, spruce, and pine forest to the pastures. The total study area of the

Himalayan and Hindukush range is 7656.91 sq. km. The categorization of the area is based on three potential thresholds; for Himalayan Monal, the highly suitable habitat is 844.4 sq. km, while the moderately suitable habitat is 2819.42 sq. km, (Table 3) and the remaining one is the not-suitable (poor) habitat predicted by the model. We also used the Cringing model to predict the hot spots, Bar Palas region of Koli Palas district and Jalkot and Kandia valley of district upper Kohistan is predicted as core zones or hot spots for Himalayan Monal (Fig 3). The Koklass pheasant predicted habitats also occupying a significant area of the study region were 611.5 sq. km, 2551.3 sq. km, and 4494.11 sq. km highly suitable, moderately suitable, and not suitable respectively. Jalkot and Kandia valley of upper Kohistan Kayal valley of lower Kohistan predicted as hot spots for Koklass pheasant predicted by the Cringing model (Fig 4). The study also quantified the potential habitat of these species from all the studying valleys of the Himalayan and Hindukush ranges. (Fig 5).

**Table 3.** Quantification of the current potential habitat based on three threshold for Himalayan Monal and Koklass Pheasant species.

| **S.No** | **Species** | **Total Area** | **Suitable habitat** | **Moderate Suitable habitat** | **Not-Suitable/Poor habitat** |
| --- | --- | --- | --- | --- | --- |
| 1 | Himalayan Monal | 7656.91sq.km. | 844.4 sq. km | 2819.42sq.km | 3993.09sq.km |
| 2 | Koklass Pheasant | 7656.91sq.km. | 611.5 sq. km | 2551.3 sq. km | 4494.11 sq. km |

**Fig 3.**
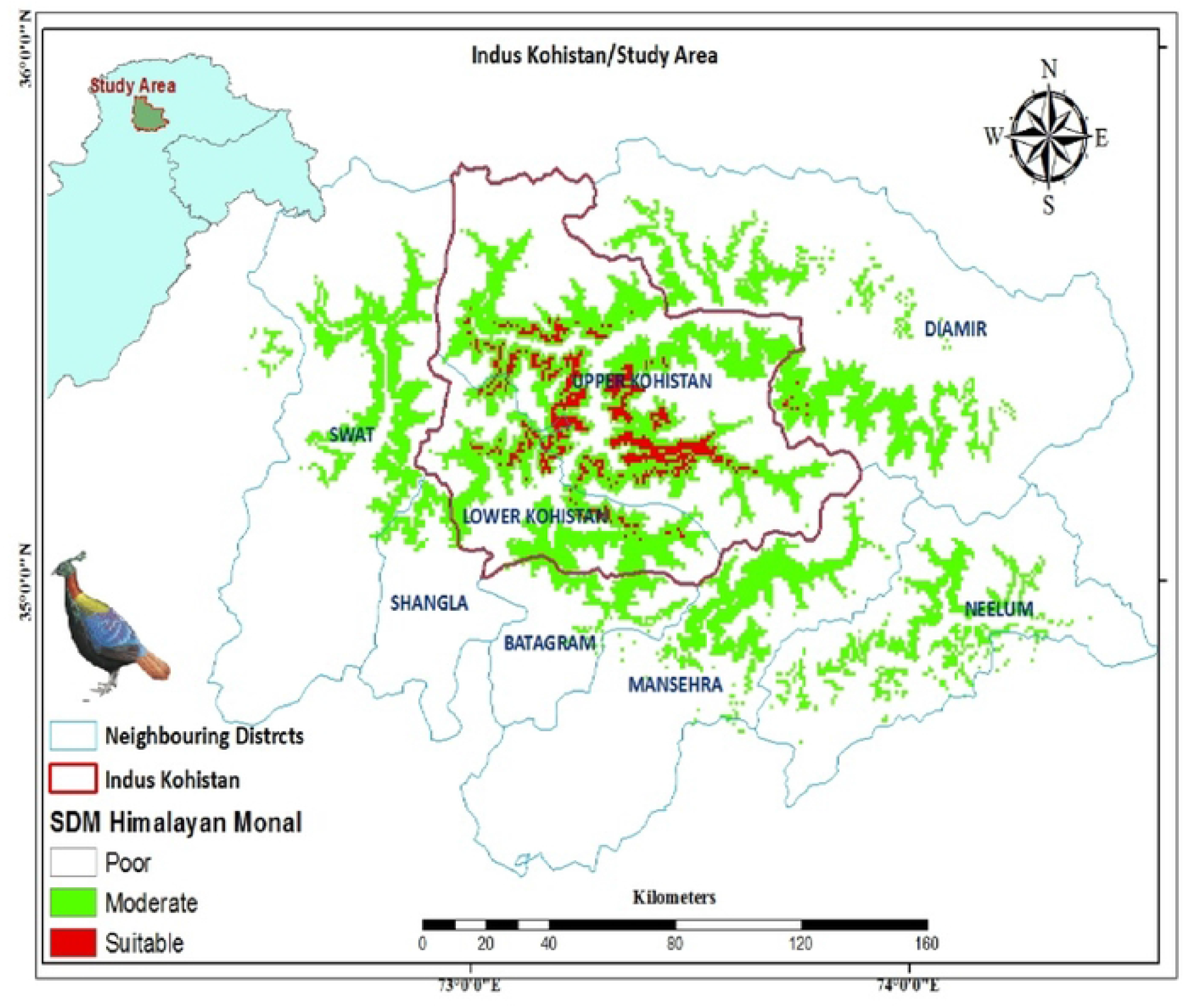
Habitat suitability of Himalayan Monal in Himalayas and Hindukush ranges special reference topheasant in Indus Kohistan, Pakistan.

**Fig 4.**
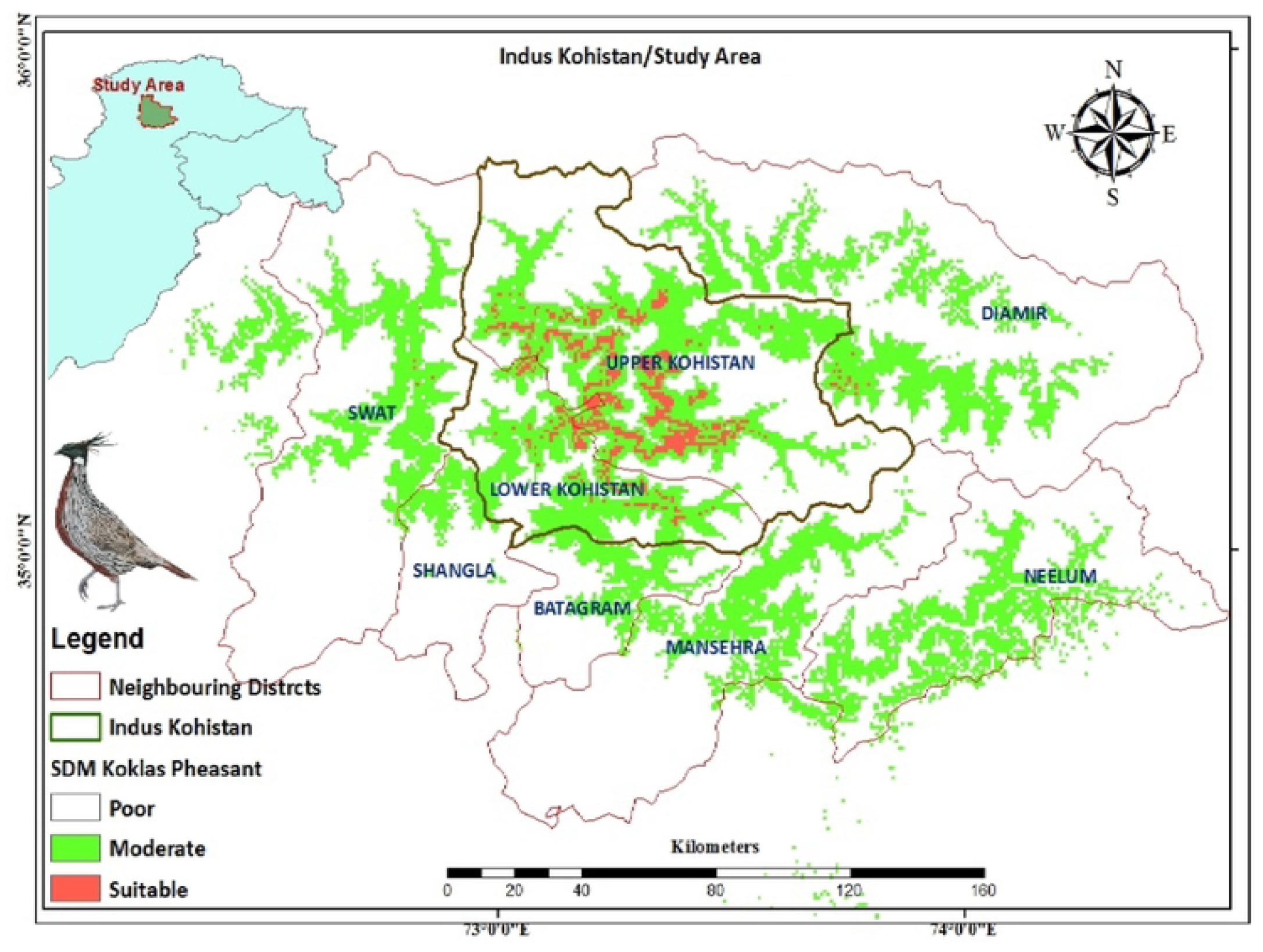
Habitat suitability of Koklass pheasant in Himalayas and Hindukush ranges special reference topheasant in Indus Kohistan, Pakistan.

**Fig 5.**
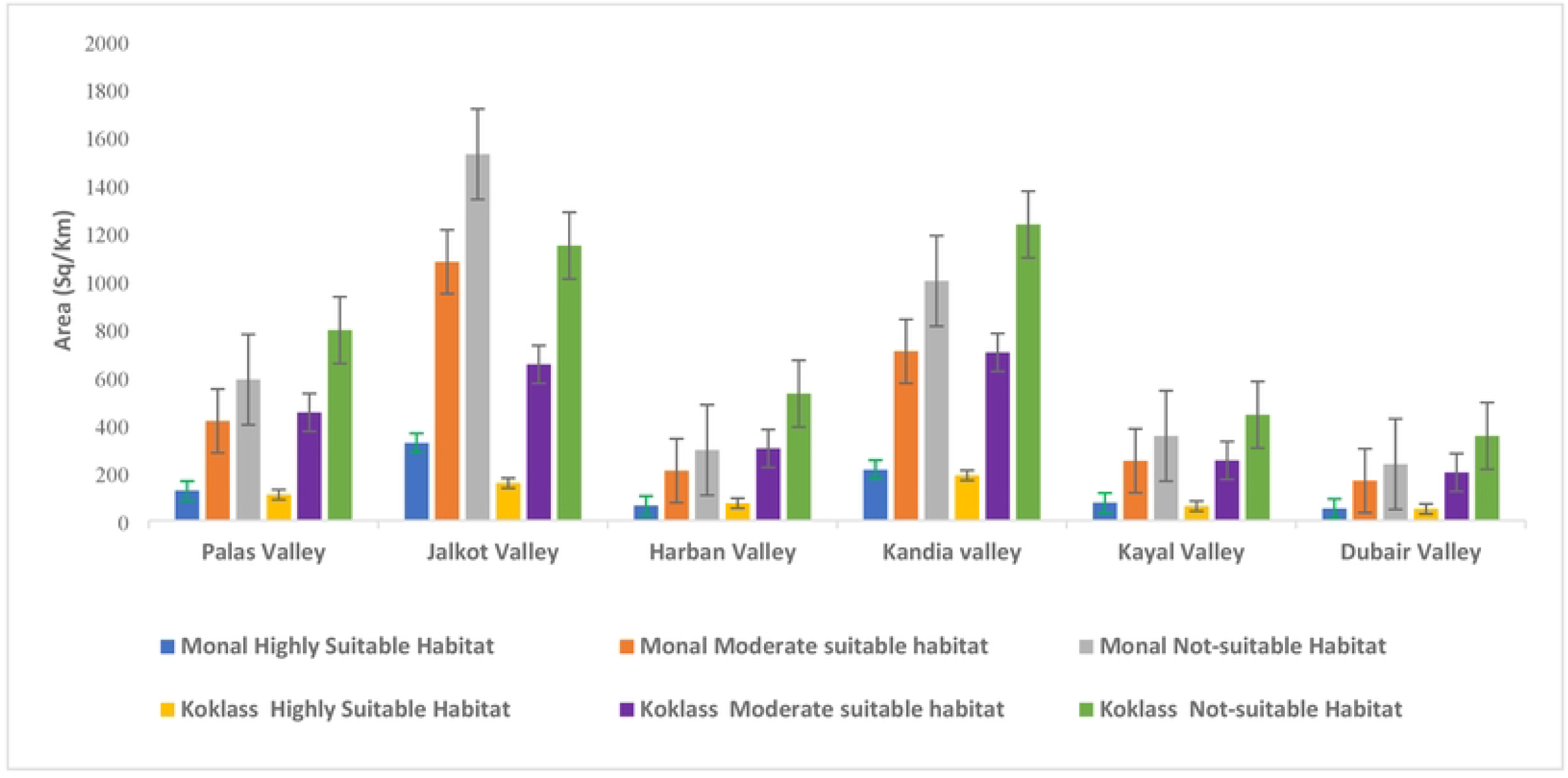
Quantification of the current potential habitat of the species is dependent on each threshold class in different valleys of the western Himalayas and Hindukush with special reference to Indus Kohistan, Pakistan.

## Discussion

The current study represents the first-ever large-scale assessment of Himalayan Monal and Koklass pheasant distribution, habitat suitability, and prediction of the hotspots in the Himalayan and Hindukush ranges in northern areas of Pakistan. The study also quantified the suitable habitat with their suitability index, the suitable and moderate suitable area is smaller than the not-suitable area and the suitable habitat is degrading rapidly due to urbanization, deforestation, and heavy construction. Our model showed considerable predictions regarding AUC value > 0.9 that may be placed among the highest in published models [31,32], and excellent values of the True Skill Statistic, corresponding to a very high predictive capacity [33].

The pheasant species show seasonal migration behavior from lower to higher altitudes and gain up to 2400m to 4500m in summer, and flashback to 1350m in winter [34,35,36]. The pheasant species prefers to inhabit and roost in the forest of *Quercus semecarpifolia, Betula utilis, Acer caecium, Juglans regia, Pinus wallichiana, Picea smithiana, Cedrus deodara* and *Abies pindrow*) [36]. The current study also revealed that the Himalayan Monal shows a greater range of seasonal migration reaching up to 4200m in summer and flashback to their winter habitat of 2000m, in hot summer the bird reaches up to alpine pastures. The Koklass pheasant also shows the seasonal migration up to 3500m in summer and back to their winter habitat of 1600m, this bird never crosses the tree line in summer. The most suitable habitats and hot spot zones are identified and predicted by Maxent and cringing model the areas where less human interference, significant forest covers, and gentle slopes are predicted as hot spots for the pheasants. The model also predicted that the areas having more human settlements, lowest and highest altitudes, pastures and open land covers have less or no suitable habitat for Koklass pheasant but the upper ranges of Himalayan Monal suitable habitat reach in pastures and open land covers. The Maxent model also predicts the suitable habitat in adjacent valleys around the study area for the specified species, in this study, the model predicted suitable habitat for both Himalayan Monal and Koklass pheasant in the adjacent valleys of other districts. The suitable habitat of neighboring districts mostly occurs in the Himalayan ranges and some are in the Hindukush ranges of Pakistan. The predicted suitable habitat of the neighbor’s district includes the Neelum District of Azad Jammu Kashmir, Mansehra, Battagram, Shangla, and Swat districts of Khyber Pakhtunkhwa, and Diamer district of Gilgit Baltistan (Fig 4).

The distribution of species in space is dependent on biotic and abiotic factors including climatic factors, and ecological factors. Understanding of these factors is vital for conservation and management purposes [37,38]. However, it is very difficult to study cryptic and shy species in inaccessible areas [39]. The climatic factors which primarily contributed to Himalayan Monal and Koklass pheasant habitat prediction, included temperature, precipitation of warmest quarter followed by annual precipitation respectively. The population of these species moves to higher altitudes in the warmest months of summer and migrate to lower altitudes in the coldest months. The perceptions of the driest months contribute maximum to their habitat prediction in Machiara National Park, Azad Jammu Kashmir followed by topographical variations like altitudes and slopes [16]. The topographical variables, altitude, slope, and distance to settlements also contributed to the habitat prediction of the species in Indus Kohistan. The slopes also contributed maximum among topographical parameters, the population mostly preferring the gentle slopes neither tough ones nor on tops of mountains. These birds are very shy and prefer to reside and roost inside a bushy forest and sometime during early dawn and dusk, appear in an open land area to searching food [13,36]. The Himalayan Monal is mostly recorded from open pastures or hillcrest during summer while in winter mostly calls from the bushes and thick forest. On the other hand, the Koklass pheasant is mostly recorded from a bushy and thick forest with 80-85% understory vegetation cover. These pheasants mostly roost during the night on lower branches of trees preferring the vintage trees. The Himalayan pheasants are very shy and the human disturbance is declining the population of the pheasants, and the natural habitat of the pheasants is being degraded [24]. The model also predicted that the parameter distance to settlements is also contributing to the habitat prediction of the species. This parameter is of prime importance for the distribution of this species, urbanization, deforestation, and heavy construction contributed the maximum to the decline in this specific area. The Himalayan Monal and Koklass pheasant are ground-dwelling and roosting in the forest of oak, deodar, alpine fir, and spruce forest with gaining altitudes higher than 3000m [36,40]. Geographical distribution of pheasants predicted by the MaxEnt model occupying the ranges from the lower regions of a sub-tropical thick forest of oaks, tropical conifer forest including deodar, fir, spruce, and pine forest to the pastures.

## Conclusion

MaxEnt model showed excellent predictive performance showed the strong prediction of the probability distribution and habitat suitability. The environmental parameters like temperature precipitation of warmest quarter contributed the maximum followed by annual precipitation to predict the habitat of pheasant species. The topographical variables, altitude, slope, distance to settlements, and the biophysical parameter mosaic land cover contributed to predicting the habitat of these species. The habitat predictions by using three thresholds for Himalayan Monal’s most suitable habitat is 844.4 sq. km while for the Koklass pheasant were 611.5 sq. km respectively. The study also predicted the hot spots by using the Cringing model; bar Palas valley, Jalkot, and Kandia valley are predicted as core zones or hot spots for Himalayan Monal, while Jalkot, Kandia, and Kayal valley are predicted as hot spots for Koklass pheasant. The study recommends that the predicted suitable habitat and valleysshould be declared as a protected area for the conservation of both pheasant species.

## Acknowledgments

Thanks to the Wildlife Department of Khyber Pakhtunkhwa for support during the fieldwork and data collection, and the Ecology Lab University of Haripur for software and analysis.

